# Proteomic and Metabolomic Profiling Nominates Druggable Targets and Biomarkers for Pulmonary Arterial Hypertension-Associated Myopathy and Exercise Intolerance

**DOI:** 10.1101/2025.03.27.644723

**Authors:** Phablo Abreu, Ryan Moon, Jenna B. Mendelson, Todd Markowski, LeeAnn Higgins, Kevin Murray, Candace Guerrero, Jeffrey Blake, Sasha Z. Prisco, Kurt W. Prins

**Affiliations:** Lillehei Heart Institute, Cardiovascular Division, Department of Medicine, University of Minnesota Medical School, Minneapolis, Minnesota, USA; Department of Integrated Biology and Physiology, University of Minnesota, Minneapolis, Minnesota, USA; Center for Metabolomics and Proteomics, Department of Biochemistry, Molecular Biology, and Biophysics, University of Minnesota, Minneapolis, Minnesota, USA

## Abstract

**Background:** Pulmonary arterial hypertension (PAH) is a rare but debilitating condition that causes exercise intolerance and ultimately death. Skeletal muscle derangements contribute to depressed exercise capacity in PAH, but the mechanisms underlying muscle dysfunction including the changes in muscle biology based on fiber type are understudied.

**Methods:** We evaluated exercise capacity, muscle histopathology, mitochondrial density, mitochondrial proteomics, and metabolomics/lipidomics of *quadriceps (*predominately fast fibers) and *soleus (*predominately slow fibers) muscles in the monocrotaline (MCT) rat model of PAH.

**Results:** MCT rats exhibited impaired exercise capacity. Surprisingly, there were divergent atrophic and metabolic remodeling in the *quadriceps* and *soleus* muscles of MCT rats. In the *quadriceps*, there was a mild atrophic response only in type II fibers. In contrast, both type I and II fibers atrophied in the *soleus*. Both muscles exhibited fibrotic infiltration, but mitochondrial density was reduced in the *quadriceps* only. Mitochondrial proteomics and tissue metabolomics/lipidomics profiling demonstrated the two muscles exhibited distinct responses as the *quadriceps* had impairments in oxidative phosphorylation/fat metabolism and storage of triacylglycerides. However, the *soleus* showed signs of proteasome deficiencies and alterations in phosphatidylcholine/phosphatidylethanolamine homeostasis. Finally, profiling of metabolites/lipids in the serum identified potential novel biomarkers of exercise intolerance in PAH including the dimethylarginine pathway, cysteine, and triacylglycerides.

**Conclusion:** Our data suggests differential cachectic and metabolic responses occur in PAH-induced myopathy. We nominate mitochondrial biogenesis and proteasome activation as potential druggable targets for PAH-myopathy.

## INTRODUCTION

Pulmonary arterial hypertension (PAH) is a rare but deadly disease that results from adverse pulmonary vascular remodeling, ultimately leading to right heart failure and death(1). One of the most commonly reported symptoms in PAH is exercise intolerance(2). Depressed exercise capability in PAH is multifactorial as there is an inability to augment cardiac output due to right ventricular systolic and diastolic impairments, insufficient ventilatory response, and skeletal muscle dysfunction(2). Thus, modulation of any one of these pathologically altered organ systems could enhance exercise capacity in PAH.

Skeletal muscles are comprised of multiple fiber types that exist on a continuum from slow to fast twitch, which allows them to perform distinct tasks(3). Each muscle type responds to work load and cachectic stimuli in specific ways, which impact that individual muscle’s function under physiological and pathological stressors(3). Previous work suggests metabolic alterations are present in PAH skeletal muscle in both rodents(4, 5) and humans(6–8), however an evaluation of fiber-type specific changes is lacking. Understanding how distinct muscles change in PAH-induced myopathy is important as we need to define targetable pathways to counteract exercise intolerance in PAH to improve quality of life and survival. At present, we do not know if different muscles or muscle types require distinct therapies to augment their function and describing how multiple muscles with different compositions alter their molecular signatures using unbiased approaches could help define therapeutic targets for PAH-associated myopathy.

Here, we examined exercise capacity, muscle histopathology, mitochondrial density, mitochondrial protein regulation, and metabolism of the *quadriceps*, a predominately fast-twitch muscle, and the *soleus*, a predominately slow-twitch muscle, in the monocrotaline (MCT) rat model of PAH. Then, we defined how the molecular and metabolic signature of each muscle was associated with exercise capacity to provide new insights and potentially nominate therapeutic approaches for PAH-associated myopathy.

## METHODS

### Animal Model

Adult male Sprague-Dawley rats (Charles River Laboratories) received a single subcutaneous injection of MCT (60 mg/kg, Sigma-Aldrich) (*n*=5) to induce PAH. Age and sex-matched rats (*n*=5) were injected with phosphate buffered saline (PBS) and served as controls. Endpoint analysis was performed 25 days post MCT injection. All animal studies adhered to the institutional and national guidelines for the care and use of laboratory animals, as specified by the University of Minnesota Institutional Animal Care and Use Committee.

### Maximal Exercise Testing

Exercise testing was performed using the same protocol as McCullough(5) on an Omnitech Treadmill equipped with metabolic chambers. One control animal did not run on the treadmill during the maximal test, and was therefore excluded from the analysis of exercise capacity.

### Cardiopulmonary Evaluation

Echocardiography and closed-chest hemodynamics quantified cardiac function and hemodynamic alterations induced by MCT as we have described(9–12). Echocardiograms were performed by KWP and hemodynamic testing was performed by JBM.

### Histological Analysis

Picrosirius red staining of formalin fixed paraffin embedded sections delineated the fibrotic area in the *quadriceps* and *soleus* muscles. Images were blindly quantified using FIJI by RM by thresholding images to highlight fibrotic areas.

### Confocal Microscopy Analysis

Paraffin-embedded skeletal muscle sections were deparaffinized and subjected to antigen retrieval using Reveal Decloaking solution(13). Sections were permeabilized with 1% Triton X-100 in PBS and then washed/blocked in 5% goat serum in PBS. Primary antibodies to myosin heavy chain slow (BA-F8, DSHB) and myosin heavy chain type II (SC-71, DSHB) were incubated at 4°C overnight. Sections were washed/blocked in 5% goat serum and stained with AlexaFluor568 conjugated secondary antibodies at 37°C for 30 minutes. Sections were washed with PBS and then mounted in ProLong Glass Antifade Mountant (ThermoFisher Scientific). Images were collected on a Zeiss LSM900 Airyscan 2.0 microscope and then cardiomyocyte cross sectional area was blindly determined by RM using FIJI software.

### Electron Microscopy

Mitochondrial density was determined by electron microscopy in the *quadriceps* and *soleus* muscles. Briefly, skeletal muscle specimens were fixed in 4% paraformaldehyde + 1% glutaraldehyde in 0.1M PBS, pH 7.2. After fixation, tissue was washed with PBS, stained with 1% osmium tetroxide, washed in H_2_O, stained in 2% uranyl acetate, washed in H_2_O, dehydrated through a graded series of ethanol and acetone and embedded in Embed 812 resin. Following a 24-hour polymerization at 60°C, 0.1 μM ultrathin sections were prepared and post-stained with lead citrate. Electron micrographs were acquired on a JEOL 1400 Plus transmission electron microscope at the Mayo Clinic as we have previously performed in cardiac muscle(14).

### Skeletal Muscle Proteomics

Mitochondrial enrichments from the *quadriceps* (*n*=5 control and *n*=5 MCT) and *soleus* muscles (*n*=4 control and *n*=4 MCT) were purified using a Mitochondrial Isolation Kit (Abcam). Samples were then subjected to TMTpro 18-plex proteomics analyses as described(13, 14). Relative protein abundances were determined using Proteome Discoverer version 3.1 software.

### Blood Lactate Testing

Immediately upon completion of the maximal exercise test, blood was collected from the tail vein and lactate concentration was determined using the Lactate Plus Measuring Meter (Nova Biomedical).

### Tissue and Serum Metabolomics/Lipidomics Analyses

Serum and muscle specimens were evaluated using the biocrates MxP® Quant 500XL kit at the University of Minnesota Center for Metabolomics and Proteomics as previously described(13, 15) using an Agilent 6495C mass spectrometry platform.

### Correlational Analysis of Proteomics/Metabolomics Data

Determined protein abundances and metabolite absolute concentrations or relative concentrations were correlated with VO_2max_ using Metaboanalyst software (https://www.metaboanalyst.ca/).

### Pathway Analysis

Proteins that were either positively or negatively associated with VO_2max_ (*p*<0.05) using Pearson correlation test were subjected to KEGG pathway analysis using ShinyGO software v0.82 (https://bioinformatics.sdstate.edu/go/).

### Statistical Analyses

Statistical tests were performed using Prism 10.3 (GraphPad Software). *p*-values were calculated using either unpaired *t*-tests and Mann-Whitney tests based on whether data were normally distribution as determined by Shapiro-Wilk Test. Graphs show the mean (if normally distributed) or median value (non-normal distribution) and all individual values. *p*-values of <0.05 were considered to indicate significance.

## Results

### MCT rats had reduced maximal exercise capacity and altered measures of systemic metabolism

To quantify exercise capacity, we determined the maximal volume of oxygen consumption (VO_2max_) of control and MCT rats using treadmill testing combined with metabolic chambers. We validated the PAH phenotype of MCT rats using echocardiogram and hemodynamics (**Supplemental Figure 1**). MCT rats exhibited a significant reduction in VO_2max_ (Control: 67.3±9.5, MCT: 29.4±8.8, *p*=0.02), time ran on treadmill (Control: 13.5±1.1, MCT: 9.0±1.1, *p*=0.03), and maximal treadmill speed tolerated (Control Median: 26.8 [25%-75%: 21.2, 26.8], MCT Median: 19.3 [25%-75%:17.3, 19.3], *p*=0.04) (**Figure 1 A-C**). In addition, blood lactate level at exhaustion was significantly higher in MCT rats (Control: 3.9±0.5 mM, MCT: 6.9±0.8 mM, p=0.02) (**Figure 1D**). The respiratory exchange ratio (RER), a measure of systemic metabolism, was greater in MCT rats at exhaustion (Control: 0.8±0.04, MCT: 1.0±0.03, *p*=0.01) (**Figure 1E**). The higher RER value in MCT rats implied these animals utilized anaerobic carbohydrate metabolism to a greater extent than controls, while the lower RER suggested controls utilized fatty acid oxidation for energy production.

**Figure 1:**
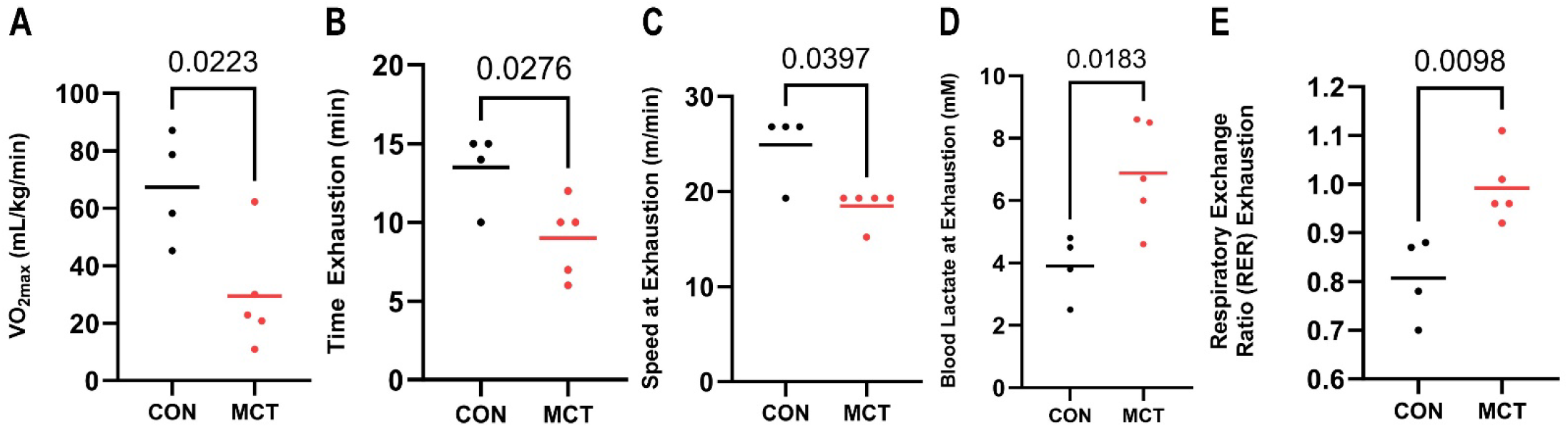
MCT Rats Exhibited Reduced Exercise Capacity and Disturbed Systemic Metabolism With Exercise. MCT rats had reduced VO_2max_ (**A**), tolerated time on treadmill (**B**), and lower peak treadmill speed (**C**) when compared to control rats. (**D**) At time of exhaustion, blood lactate concentration was higher in MCT rats. (**E**) At VO_2max_, the respiratory exchange ratio (RER) was elevated in MCT rats. *p*-values were calculated using either an unpaired *t*-test or Mann-Whitney U-test.

### The quadriceps and soleus exhibited divergent cachectic but comparable fibrotic responses

Next, we investigated how MCT-PAH altered the histological phenotype in *quadriceps* and *soleus* muscles. Surprisingly, we observed divergent phenotypes in the *quadriceps* and *soleus*. In the *quadriceps*, there was a shift in the fiber size distribution with a mild atrophic response in fast-twitch type II fibers (Control Median: 929 μm^2^ [25%-75%: 715 μm^2^, 1204 μm^2^], MCT Median: 871 μm^2^ [25%-75%: 676 μm^2^, 1175 μm^2^], *p*=0.0005), but a significant increase in type I fiber cross sectional area (Control Median: 1825 μm^2^ [25%-75%: 1416 μm^2^, 2242 μm^2^], MCT Median: 2157 μm^2^ [25%-75%: 1633 μm^2^, 2945 μm^2^], *p*<0.0001) (**Figure 2 A-C**). In contrast, in the soleus, there was a consistent reduction in fiber size in both type I (Control Median: 2524 μm^2^ [25%-75%: 2092 μm^2^, 3005 μm^2^], MCT Median: 2006 μm^2^ [25%-75%: 2006 μm^2^, 2340 μm^2^], *p*<0.0001) and II fibers (Control Median: 2239 μm^2^ [25%-75%: 1922 μm^2^, 2528 μm^2^], MCT Median: 1634 μm^2^ [25%-75%: 1333 μm^2^, 1970 μm^2^], *p*<0.0001) (**Figure 2 D-F**). Finally, we found heightened fibrosis in both the *quadriceps* (Control: 2.9±0.2%, MCT: 10.5±0.6%, *p*<0.0001) and *soleus* (Control: 2.6±0.2%, MCT: 9.7±0.7%, *p*<0.0001) (**Figure 2 D-F**).

**Figure 2:**
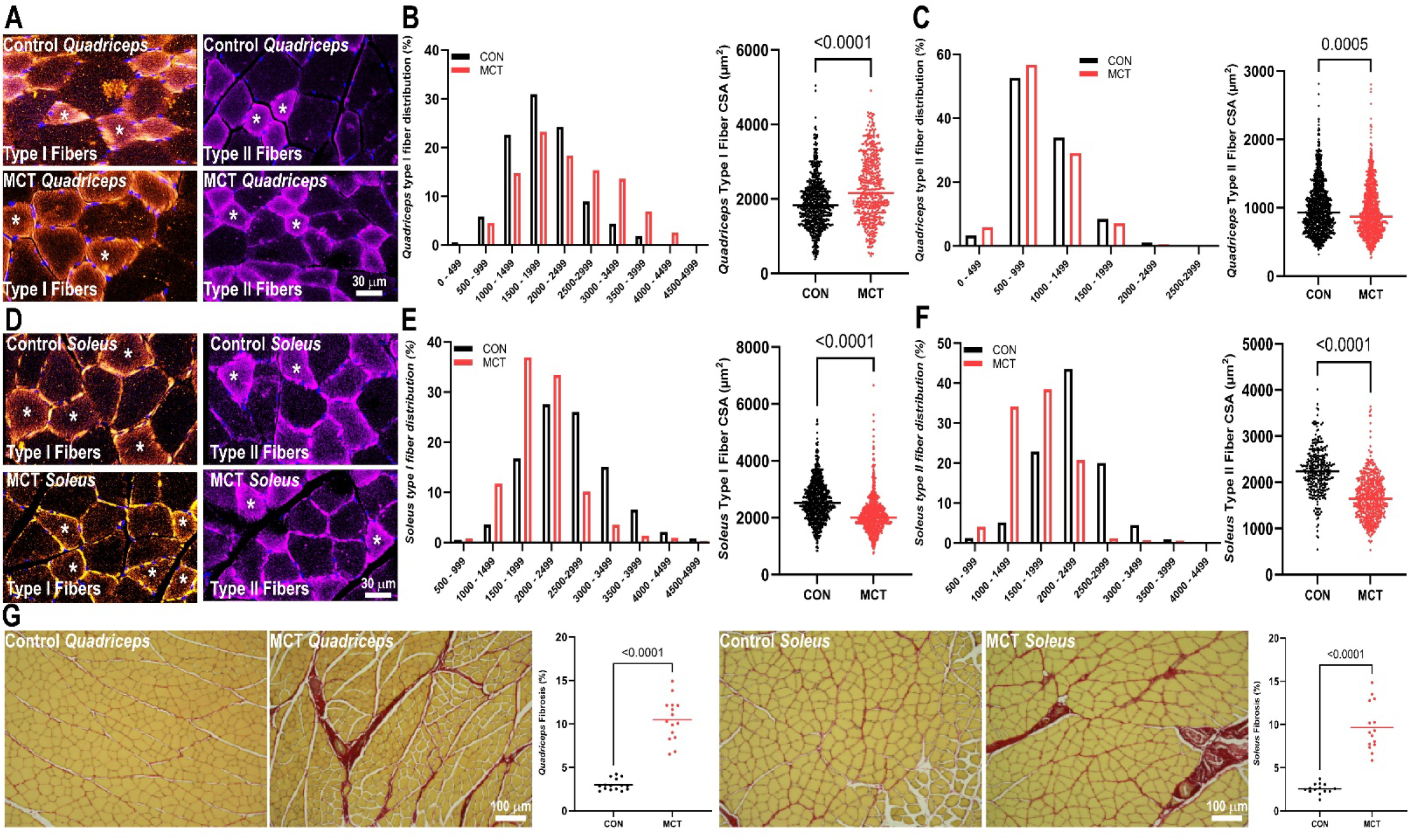
The *Quadriceps* and *Soleus* Muscles Exhibited Divergent Atrophic Phenotypes but Conserved Fibrotic Responses. Representative confocal micrographs depicting *quadriceps* sections from control and MCT rats stained with antibody to delineate type I (orange) and type II (purple) fibers (**A**). (**B**) Distribution of type I fiber sizes and quantification of cross-sectional area in *quadriceps* sections. (C) Quantification of type II size distribution and type II cross-sectional area in *quadriceps* sections. (D) Representative confocal micrographs depicting *soleus* sections from control and MCT rats stained with antibody to delineate type I (orange) and type II (purple) fibers. (E) Distribution of type I fiber size and quantification of cross-sectional area in *soleus* sections. (F) Size distribution and quantification of cross-sectional area of type II fibers in *soleus* sections. (G) Representative picrosirius stained section and quantification of fibrosis in the *quadriceps* and *soleus. p*-values were calculated using either an unpaired *t*-test or Mann-Whitney U-test.

### Mitochondrial density and protein regulation were distinctly altered in the quadriceps and soleus muscles

We next evaluated mitochondrial density and protein regulation in the *quadriceps* and *soleus*. Electron microscopy analysis revealed a reduction in mitochondrial content in the *quadriceps* (Control Median: 3.3% [25%-75%: 1.3%, 4.0], MCT Median: 1.3% [25%-75%: 1.0%, 1.8%], *p*=0.01), but an increase in the *soleus* (Control: 5.3±0.6%, MCT: 8.6±0.7%, *p*=0.001) (**Figure 3A-D**).

**Figure 3:**
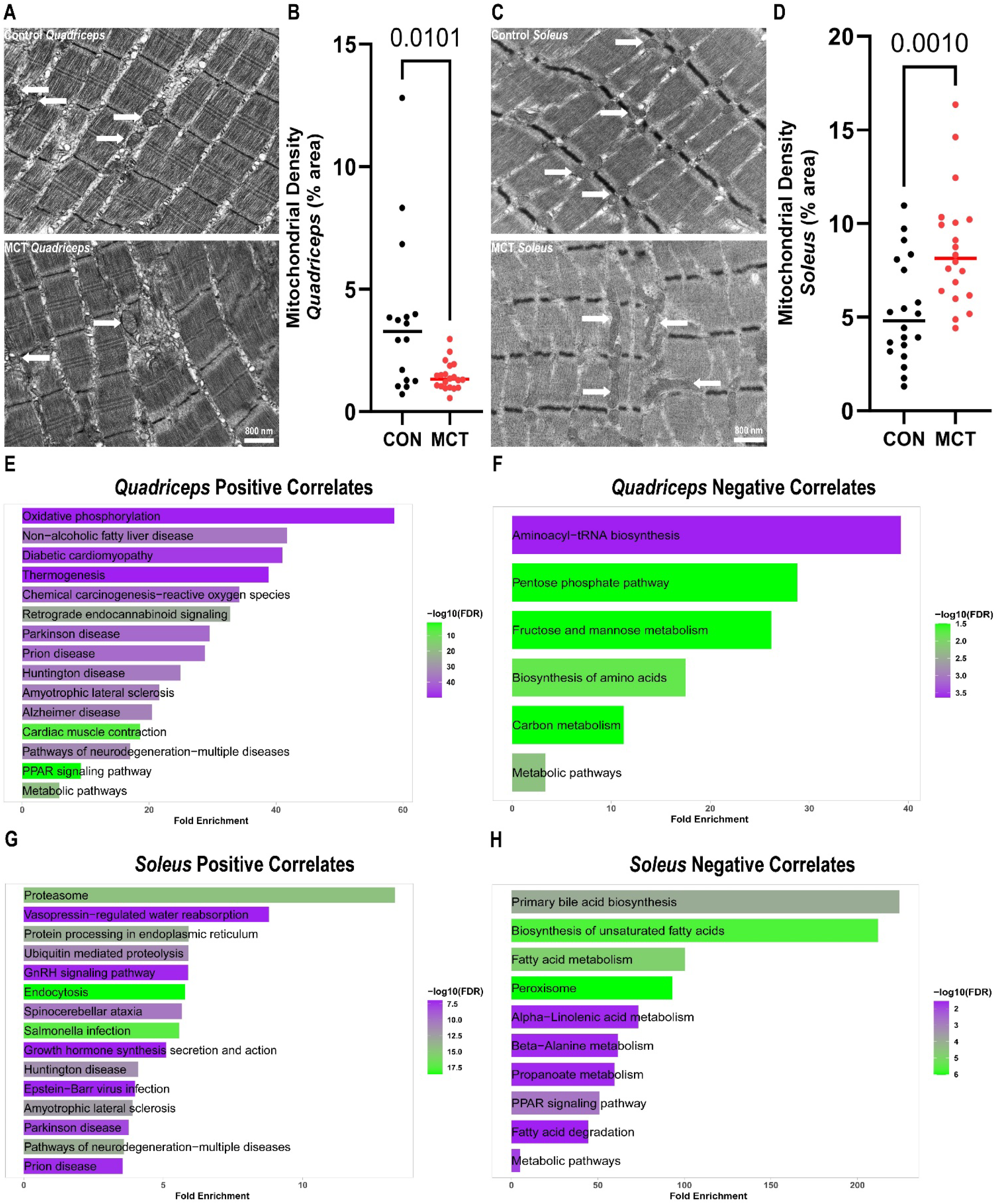
Differential Mitochondrial Density and Proteomic Responses in the *Quadriceps* and *Soleus* of MCT Rats. (**A**) Representative electron micrographs from the *quadriceps* muscles and quantification of mitochondrial density (**B**). (**C**) Electron micrographs of *soleus* specimens and quantification of mitochondrial density (**D**). (**E**) Pathway analysis of proteins positively associated with VO_2max_ in the *quadriceps*. (**F**) Pathway analysis of proteins negatively associated with VO_2max_ in the *quadriceps*. (**G**) Pathway analysis of proteins positively associated with VO_2max_ in *soleus*. (**H**) Pathway analysis of proteins negatively associated with VO_2max_ in *soleus. p*-values were calculated using either an unpaired *t*-test or Mann-Whitney U-test.

Then, we used proteomics analysis to delineate mitochondrial protein regulation. We quantified the abundances of >2,900 proteins in *quadriceps* and *soleus* mitochondrial enrichments using an unbiased proteomics assessment. We used correlational analysis to define which proteins and protein pathways were significantly associated with VO_2max_. In the *quadriceps*, 91 proteins were positively associated and 50 proteins were negatively associated with VO_2max_. Pathway analysis of proteins positively associated with VO_2max_ in the *quadriceps* identified pathways associated with electron transport chain function (oxidative phosphorylation) and fatty acid metabolism (non-alcoholic fatty liver disease, diabetic cardiomyopathy, PPAR signaling pathway) (**Figure 3E**). On the other hand, pathways enriched with proteins negatively associated with VO_2max_ included those for carbohydrate and amino acid homeostasis/metabolism (**Figure 3E**). Thus, these data suggested that in the MCT *quadriceps* fatty acid/oxidative phosphorylation was impaired and carbohydrate/amino acid metabolism pathways were activated.

In the *soleus*, there were 636 proteins positively correlated, and 40 proteins negatively correlated with VO_2max_. Pathway analysis suggested protein homeostasis pathways including the proteasome, ubiquitin mediated proteolysis, and protein processing in the endoplasmic reticulum were important for *soleus*-related exercise capacity (**Figure 3F**). In contrast, pathways associated with fatty acid metabolism including the peroxisome, fatty acid metabolism, PPAR signaling, and metabolic pathways were inversely associated with VO_2max_ (**Figure 3F**). These data implied protein processing/homeostasis was disrupted in the MCT *soleus*. However, fatty acid metabolizing pathways were not suppressed and even slightly increased in the MCT *soleus*, which highlighted a divergent response between the two muscles.

### Metabolomics/Lipidomics Revealed Unique Metabolic Responses in Quadriceps and Soleus Muscles

Next, we implemented a combined metabolomic/lipidomic analysis to delineate how alterations in mitochondrial content and protein regulation impacted muscle metabolism. We quantified >1000 metabolites/lipids or metabolite ratios in each muscle and performed correlational analysis to delineate which metabolites were most associated with exercise capacity. In the *quadriceps*, there were 121 metabolites/lipid species that were significantly associated with VO_2max_, of which 116 were positively correlated and 5 were negatively correlated. The strongest association was between triacylglycerides and VO_2max_ (**Figure 4A and B**), and nearly all triacylglycerides were reduced in the MCT *quadriceps* when compared to controls (**Figure 4C**).

**Figure 4:**
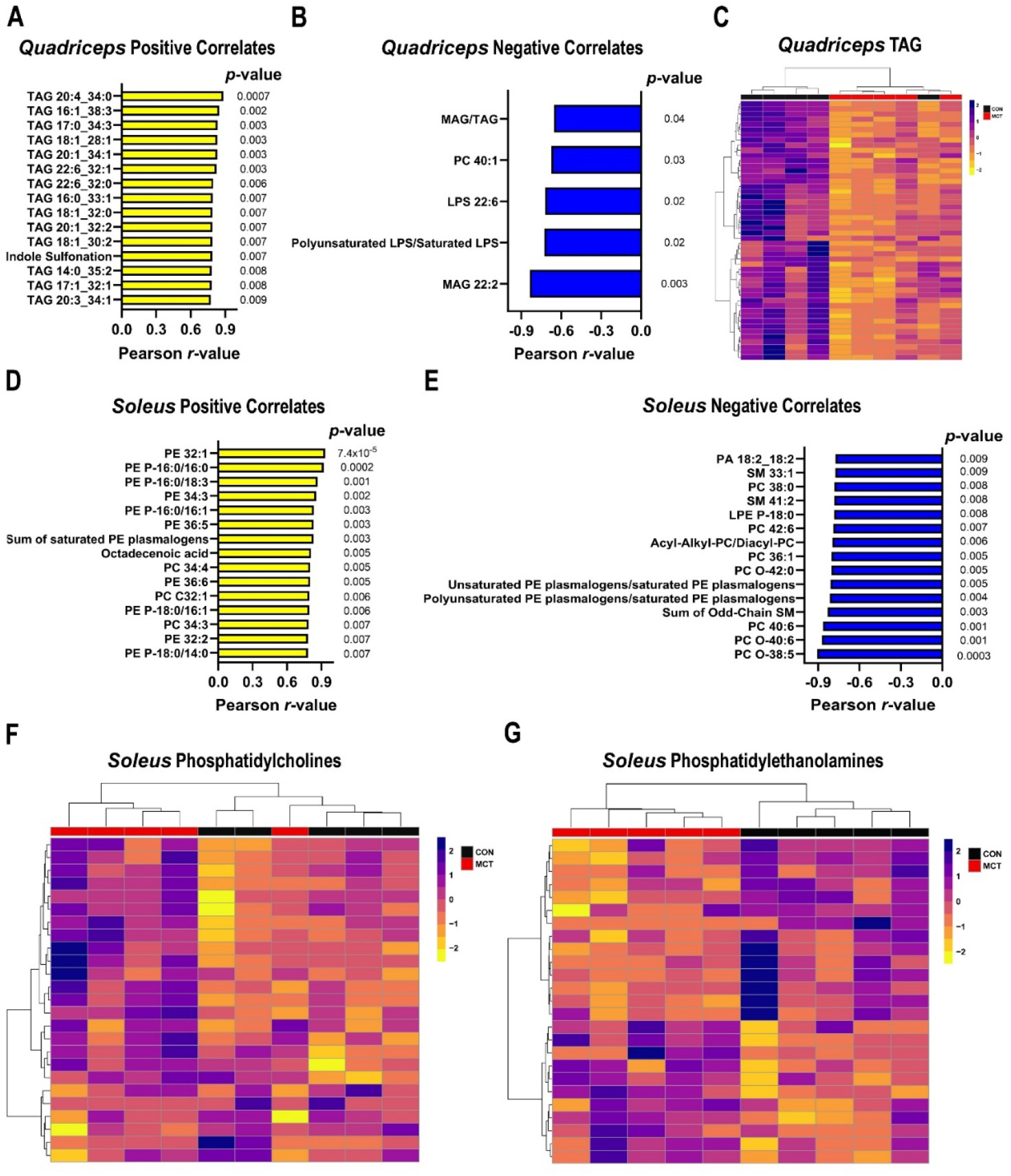
Correlational Analysis Identified Distinct Metabolic and Lipidomic Remodeling in the *Quadriceps* and the *Soleus*. Depiction of metabolites from *quadriceps* samples positively (**A**) or negatively (**B**) associated with VO_2max_. (**C**) Hierarchical cluster analysis of triacylglycerides (TAG) in *quadriceps* specimens. Depiction of metabolites from *soleus* samples positively (**D**) or negatively (**E**) associated with VO_2max_. Hierarchical cluster analysis of phosphatidylcholines (**F**) or phosphatidylethanolamines (**G**) in *soleus* specimens.

In the *soleus*, there were 103 metabolites significantly correlated with VO_2max_, with 57 positively associated and 46 negatively associated. The strongest signal was in lipid biology as numerous phosphatidylethanolamines were correlated with improved exercise performance (**Figure 4D**) while several phosphatidylcholines were inversely related to exercise capability (**Figure 4E**). Overall, several phosphatidylcholines were elevated in MCT animals (**Figure 4F**) while many phosphatidylethanolamines were reduced (**Figure 4G**).

### Serum Metabolomics/Lipidomics Suggested Systemic Metabolic Derangements Were Present and Nominated Potential Biomarkers for PAH-Associated Exercise Intolerance

Finally, we performed metabolomics/lipidomics analysis of serum samples to both gain insights into the effects of systemic metabolism on exercise capacity and nominate potential biomarkers for exercise intolerance. Overall, the serum metabolomics signature was distinct when the two groups were compared (**Figure 5A**). In total, 70 metabolites/lipids/pathways were positively associated with VO_2max_ and 30 metabolites/lipids/pathways were negatively associated with VO_2max_. We found serum triacylglycerides, cysteine homeostasis markers, and measures of dimethylarginine metabolism were the most robust readouts of exercise performance (**Figure 5 B and C**).

**Figure 5:**
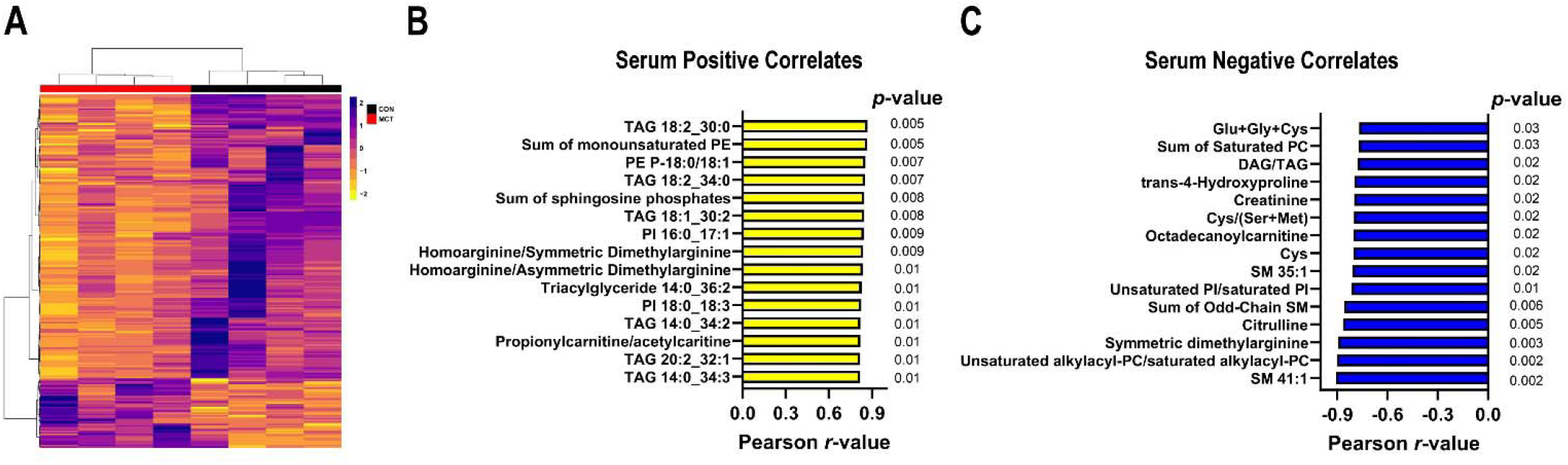
Serum Metabolomic/Lipidomic Profiling Identified Altered Systemic Metabolism and Defined Biomarkers of Exercise Capacity in MCT rats. (**A**) Hierarchical cluster analysis depicted differences in circulating metabolites in control (*n*=4) and MCT (*n*=4) rats. (**B**) 15 metabolites most strongly and positively associated with VO_2max_. (**C**) 15 metabolites most strongly and negatively correlated with VO_2max_.

## Discussion

In this study, we use proteomics and metabolomics to understand the molecular landscape of PAH myopathy and define how exercise capacity is associated with muscle remodeling. We show the *quadriceps* and *soleus* have distinct morphological and molecular responses to MCT-induced PAH. The cachectic response is more robust in the *soleus* but both muscles exhibit fibrosis. The *quadriceps* exhibits a mitochondrial biogenesis defect defined by reductions in electron transport and fatty acid metabolism proteins and a loss of triacylglycerides stores. In contrast, the *soleus* displays an impairment in proteasome regulation and imbalances of phosphatidylcholine and phosphatidylethanolamine homeostasis. The serum metabolome is distinct between the two groups and serum cysteine metabolites, dimethylarginine pathway intermediates, and triacylglycerides predict VO_2max_. Thus, we provide additional insights into the mechanisms underlying preclinical PAH myopathy and define biomarkers of exercise intolerance (**Figure 6**), which will hopefully be useful as we continue to explore new methods to enhance exercise capacity in PAH.

**Figure 6:**
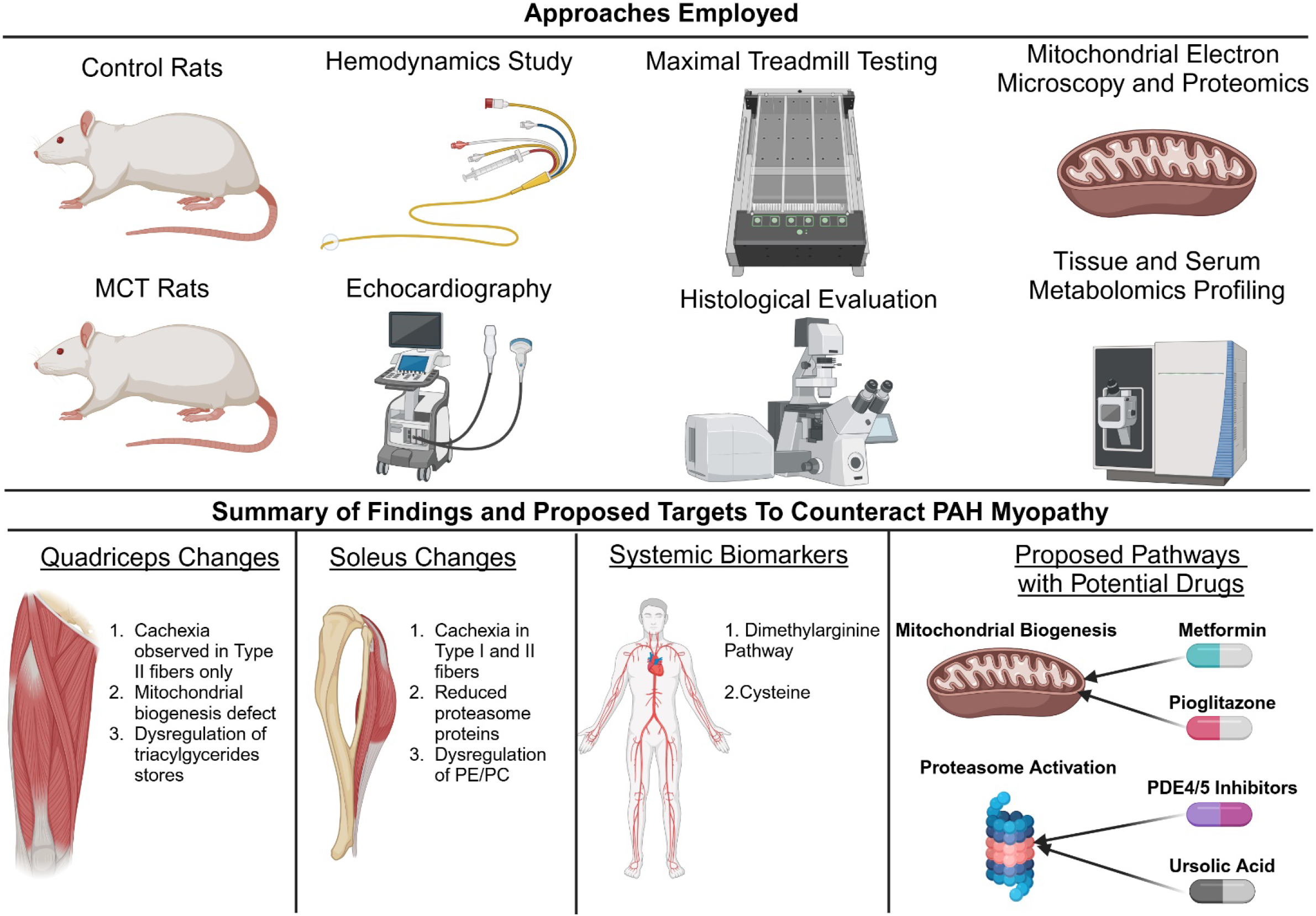
Central Figure.

Although we observed disparate proteomic and metabolomic alterations in the *quadriceps* and *soleus*, one consistent finding is that type II fibers atrophy in both muscles. Multiple studies show type II fibers more rapidly atrophy as compared to type I fibers(16), so our data is consistent with those. Our data parallel the data from the severe Fisher rat model of Sugen-hypoxia PAH, where both muscle types atrophy in the *soleus*(4). However, the divergent response of type I fibers in each muscle will need to be evaluated in greater detail in future studies. Because both muscles have significant fibrotic infiltration (**Figure 2**), it is less likely that fibrotic signaling is a driver of these distinct phenotypes. Alterations in mitochondrial function are distinct between the *quadriceps* and *soleus* based on our proteomics and metabolomics analyses, and this is congruent with other data as mitochondrial dysfunction is not consistently observed in atrophic tissue when comparing the responses of different muscle groups(17). Further studies are needed to delineate which pathways modulate the cachectic response as metabolic remodeling and fibrosis do not appear to be drivers of cachexia in the *soleus*.

We observed downregulation of electron transport chain/oxidative phosphorylation proteins in the *quadriceps* but not the *soleus* muscle of MCT rats. This is paired with another divergent response: a reduction in mitochondrial density in the *quadriceps* but an increase in the *soleus*. These data correspond well with previously published reports using both rodent and human tissue. First, in human PAH *quadriceps* samples, electron transport chain proteins are reduced as compared to healthy controls(8). Next, there is no evidence of electron transport chain protein dysregulation or protein dysfunction in the *soleus* of the Sugen-hypoxia model of PAH(4). Thus, our data and that from other investigators suggest approaches that modulate mitochondrial biogenesis and oxidative function may confer more benefits in the *quadriceps* than the *soleus*.

Using serum metabolomics profiling, we nominate the dimethylarginine pathway as an important marker of exercise intolerance in preclinical PAH. The dimethylarginine pathway is a readout of protein catabolism as dimethylarginines are found after arginine in proteins are modified and ultimately released when proteins are degraded(18). Dimethylarginine is found in two forms: symmetric (SDMA) and asymmetric (ADMA)(18). ADMA is already implicated in PAH as persistent elevations of ADMA are associated with poor outcomes(19). Unlike ADMA, which suppresses endothelial nitric oxide signaling(20), SDMA does not have clear endothelial compromising activity(18). However, SDMA is renally secreted and numerous studies demonstrate SDMA is a robust marker of renal function(21). Perhaps renal dysfunction contributes to exercise intolerance in our rodent studies and that is why SDMA is strongly associated with depressed exercise capability. SDMA may be a metabolite worth evaluating in larger, human PAH cohorts, and specifically probing its link to both exercise capacity and renal compromise.

Our data suggests that another potential biomarker of exercise intolerance in PAH is cysteine as higher serum levels of cysteine itself, or in combination with other amino acids, is associated with lower VO_2max_ (**Figure 5**). There is precedence for alterations in cysteine metabolism in pulmonary hypertension as serum cysteine predicts pulmonary artery pressure in a multivariate analysis and every 1% increase in serum cysteine is paired with a 16% increase in pulmonary artery systolic pressure(22). Furthermore, dysregulation of the cysteine pathway is associated with reduced six-minute walk test and survival in a metabolomics analysis of 117 human PAH patients(23). Serum cysteine levels in larger and multi-centered cohorts should be evaluated to determine if cysteine associates with exercise capacity and survival in PAH.

Taking our data in total, we nominate two distinct ways to counteract PAH myopathy, and there are already Food and Drug Administration approved pharmaceuticals that could be evaluated for this hypothesis (**Figure 6**). First, our data suggest approaches to promote mitochondrial biogenesis could be beneficial for the *quadriceps*. One already available drug that promotes mitochondrial biogenesis is metformin(24), and there are data that metformin counteracts PAH development and RV dysfunction in animal studies(25). Moreover, metformin is safe in PAH patients without diabetes, but in this clinical trial, metformin does not improve six-minute walk distance(26). This may be due to the fact that these patients exhibited minimal compromise with a baseline six-minute walk distance of 434 meters(26), so they may not have had significant myopathy. Another approach to promote mitochondrial biogenesis is signaling through the peroxisome proliferator-activated receptor gamma (PPARγ) pathway(27). Again, there are preclinical data supporting the use of PPARγ agonist for counteracting PAH pathobiology and right ventricular failure(28). However, its effect on skeletal muscle structure and function is not well defined, and this may be important to consider if this class of medications are evaluated in PAH as they could offer additional skeletal-muscle enhancing effects.

Finally, stimulation of proteasome activity may counteract *soleus* cachexia to improve exercise capacity. This finding was counterintuitive as proteasome activation promotes muscle atrophy in disuse situations, which suggests the *soleus* cachexia in PAH is not simply due to reduced activity. However, it is in line with other data showing knockout of key proteasome subunits in skeletal muscle promotes muscle dysfunction and wasting(29). Interestingly, there are small molecule proteasome activators that could be used to test this hypothesis. First and maybe directly relevant to PAH, activation of the proteasome occurs with phosphodiesterase inhibition(30). Phosphodiesterase inhibition is one of the cornerstone therapies for PAH(1), and these data raise the possibility that some of the beneficial effects of phosphodiesterase inhibition on exercise capacity are proteasome activation in skeletal muscle. In addition, the small molecule, ursolic acid, is a proteasome activator(31, 32) that could be evaluated for PAH myopathy. Ursolic acid counteracts PAH in rodents(33) suggesting it may also have dual effects in muscle and in the lung. In total, these data suggest augmenting proteasome activity could both counteract PAH development/progression and skeletal muscle atrophy. Consistent with this hypothesis, the proteasome inhibitor, carfilzomib is linked to PAH development(34). Further evaluation of proteasome regulators should be considered to more thoroughly evaluate this hypothesis.

## Limitations

Our study has important limitations that we acknowledge. First, we only use male animals. One control animal did not complete the maximal treadmill test which reduced some of our power for analyzing metabolic and proteomic associations with exercise capacity. In addition, caloric consumption can impact metabolomics/lipidomics findings in both the tissue and in the serum, and we previously demonstrated MCT rats consume less food than controls(14). This may contribute to the reduction of serum triacylglycerides in MCT rats, which is not what is consistently observed in PAH patients(35). We therefore do not believe serum triacylglycerides are an appropriate measure of exercise capacity in human PAH.

## Acknowledgements

We thank Trace Christensen and the Mayo Microscopy and Cell Analysis Core for their assistance in collecting electron micrographs.

## REFERENCES

1. Thenappan T, Ormiston ML, Ryan JJ, Archer SL: Pulmonary arterial hypertension: pathogenesis and clinical management. BMJ 2018;360:j5492.

2. Malenfant S, Lebret M, Breton-Gagnon É, et al.: Exercise intolerance in pulmonary arterial hypertension: insight into central and peripheral pathophysiological mechanisms. Eur Respir Rev 2021;30.

3. Mukund K, Subramaniam S: Skeletal muscle: A review of molecular structure and function, in health and disease. Wiley Interdiscip Rev Syst Biol Med 2020;12:e1462.

4. Zhang P, Da Silva Goncalves Bos D, Vang A, et al.: Reduced exercise capacity occurs before intrinsic skeletal muscle dysfunction in experimental rat models of pulmonary hypertension. Pulm Circ 2024;14:e12358.

5. McCullough DJ, Kue N, Mancini T, Vang A, Clements RT, Choudhary G: Endurance exercise training in pulmonary hypertension increases skeletal muscle electron transport chain supercomplex assembly. Pulm Circ 2020;10:2045894020925762.

6. Menezes TCF, Lee MH, Fonseca Balladares DC, et al.: Skeletal Muscle Pathology in Pulmonary Arterial Hypertension and Its Contribution to Exercise Intolerance. J Am Heart Assoc 2025;14:e036952.

7. Potus F, Malenfant S, Graydon C, et al.: Impaired angiogenesis and peripheral muscle microcirculation loss contribute to exercise intolerance in pulmonary arterial hypertension. Am J Respir Crit Care Med 2014;190:318–28.

8. Malenfant S, Potus F, Fournier F, et al.: Skeletal muscle proteomic signature and metabolic impairment in pulmonary hypertension. J Mol Med (Berl) 2015;93:573–84.

9. Prisco SZ, Eklund M, Raveendran R, Thenappan T, Prins KW: With No Lysine Kinase 1 Promotes Metabolic Derangements and RV Dysfunction in Pulmonary Arterial Hypertension. JACC Basic Transl Sci 2021;6:834–50.

10. Kazmirczak F, Vogel NT, Prisco SZ, et al.: Ferroptosis-Mediated Inflammation Promotes Pulmonary Hypertension. Circ Res 2024;135:1067–83.

11. Prisco SZ, Hartweck LM, Kazmirczak F, et al.: Junctophilin-2 Regulates Mitochondrial Metabolism. Circulation 2024;150:657–60.

12. Prisco SZ, Hartweck LM, Rose L, et al.: Inflammatory Glycoprotein 130 Signaling Links Changes in Microtubules and Junctophilin-2 to Altered Mitochondrial Metabolism and Right Ventricular Contractility. Circ Heart Fail 2022;15:e008574.

13. Kazmirczak F, Vogel NT, Prisco SZ, et al.: Ferroptosis Mediated Inflammation Promotes Pulmonary Hypertension. Circ Res 2024.

14. Kazmirczak F, Hartweck LM, Vogel NT, et al.: Intermittent Fasting Activates AMP-Kinase to Restructure Right Ventricular Lipid Metabolism and Microtubules. JACC Basic Transl Sci 2023;8:239–54.

15. Kazmirczak F, Moon R, Vogel NT, et al.: Ferroptosis Inhibition Combats Metabolic Derangements and Improves Cardiac Function in Pulmonary Artery Banded Pigs. Am J Respir Crit Care Med 2024.

16. Wang Y, Pessin JE: Mechanisms for fiber-type specificity of skeletal muscle atrophy. Curr Opin Clin Nutr Metab Care 2013;16:243–50.

17. Picard M, Ritchie D, Thomas MM, Wright KJ, Hepple RT: Alterations in intrinsic mitochondrial function with aging are fiber type-specific and do not explain differential atrophy between muscles. Aging Cell 2011;10:1047–55.

18. Tain YL, Hsu CN: Toxic Dimethylarginines: Asymmetric Dimethylarginine (ADMA) and Symmetric Dimethylarginine (SDMA). Toxins (Basel) 2017;9.

19. Shafran I, Probst V, Panzenböck A, et al.: Asymmetric Dimethylarginine and NT-proBNP Levels Provide Synergistic Information in Pulmonary Arterial Hypertension. JACC Heart Fail 2024;12:1089–97.

20. Pope AJ, Karrupiah K, Kearns PN, Xia Y, Cardounel AJ: Role of dimethylarginine dimethylaminohydrolases in the regulation of endothelial nitric oxide production. J Biol Chem 2009;284:35338–47.

21. Oliva-Damaso E, Oliva-Damaso N, Rodriguez-Esparragon F, et al.: Asymmetric (ADMA) and Symmetric (SDMA) Dimethylarginines in Chronic Kidney Disease: A Clinical Approach. Int J Mol Sci 2019;20.

22. Ghasemzadeh N, Patel RS, Eapen DJ, et al.: Oxidative stress is associated with increased pulmonary artery systolic pressure in humans. Hypertension 2014;63:1270–5.

23. Pi H, Xia L, Ralph DD, et al.: Metabolomic Signatures Associated With Pulmonary Arterial Hypertension Outcomes. Circ Res 2023;132:254–66.

24. Karise I, Bargut TC, Del Sol M, Aguila MB, Mandarim-de-Lacerda CA: Metformin enhances mitochondrial biogenesis and thermogenesis in brown adipocytes of mice. Biomed Pharmacother 2019;111:1156–65.

25. Prins KW, Thenappan T, Weir EK, Kalra R, Pritzker M, Archer SL: Repurposing Medications for Treatment of Pulmonary Arterial Hypertension: What’s Old Is New Again. J Am Heart Assoc 2019;8:e011343.

26. Brittain EL, Niswender K, Agrawal V, et al.: Mechanistic Phase II Clinical Trial of Metformin in Pulmonary Arterial Hypertension. J Am Heart Assoc 2020;9:e018349.

27. Corona JC, Duchen MR: PPARγ as a therapeutic target to rescue mitochondrial function in neurological disease. Free Radic Biol Med 2016;100:153–63.

28. Hansmann G, Calvier L, Risbano MG, Chan SY: Activation of the Metabolic Master Regulator PPARγ: A Potential PIOneering Therapy for Pulmonary Arterial Hypertension. Am J Respir Cell Mol Biol 2020;62:143–56.

29. Kitajima Y, Tashiro Y, Suzuki N, et al.: Proteasome dysfunction induces muscle growth defects and protein aggregation. J Cell Sci 2014;127:5204–17.

30. VerPlank JJS, Tyrkalska SD, Fleming A, Rubinsztein DC, Goldberg AL: cGMP via PKG activates 26S proteasomes and enhances degradation of proteins, including ones that cause neurodegenerative diseases. Proc Natl Acad Sci U S A 2020;117:14220–30.

31. Coleman RA, Trader DJ: Development and Application of a Sensitive Peptide Reporter to Discover 20S Proteasome Stimulators. ACS Comb Sci 2018;20:269–76.

32. Wang N, Wang E, Wang R, et al.: Ursolic acid ameliorates amyloid β-induced pathological symptoms in Caenorhabditis elegans by activating the proteasome. Neurotoxicology 2022;88:231–40.

33. Gao X, Zhang Z, Li X, et al.: Ursolic Acid Improves Monocrotaline-Induced Right Ventricular Remodeling by Regulating Metabolism. J Cardiovasc Pharmacol 2020;75:545–55.

34. Grynblat J, Khouri C, Hlavaty A, et al.: Characteristics and outcomes of patients developing pulmonary hypertension associated with proteasome inhibitors. Eur Respir J 2024;63.

35. Hemnes AR, Luther JM, Rhodes CJ, et al.: Human PAH is characterized by a pattern of lipid-related insulin resistance. JCI Insight 2019;4.

