## Supplemental Data for "Proteomic and Metabolomic Profiling Nominates Druggable Targets and Biomarkers for Pulmonary Arterial Hypertension-Associated Myopathy and Exercise Intolerance"

Supplemental Figure 1: Hemodynamics and echocardiography validate the PAH phenotype of MCT rats. (A) MCT rats had significantly higher right ventricular systolic pressure (RVSP) as compared to control (CON: 30±2 mm Hg, MCT: 68±6 mm Hg). (B) Pulmonary artery acceleration time was reduced in MCT rats (CON: 35±1 ms, MCT: 17±1 ms). Tricuspid annular plane systolic excursion (TAPSE) was reduced in MCT rats (CON: 2.5±0.1 mm, MCT: 1.6±0.1 mm). *p*-values determined by unpaired *t*-test or unpaired *t*-test with Welch correction (A).

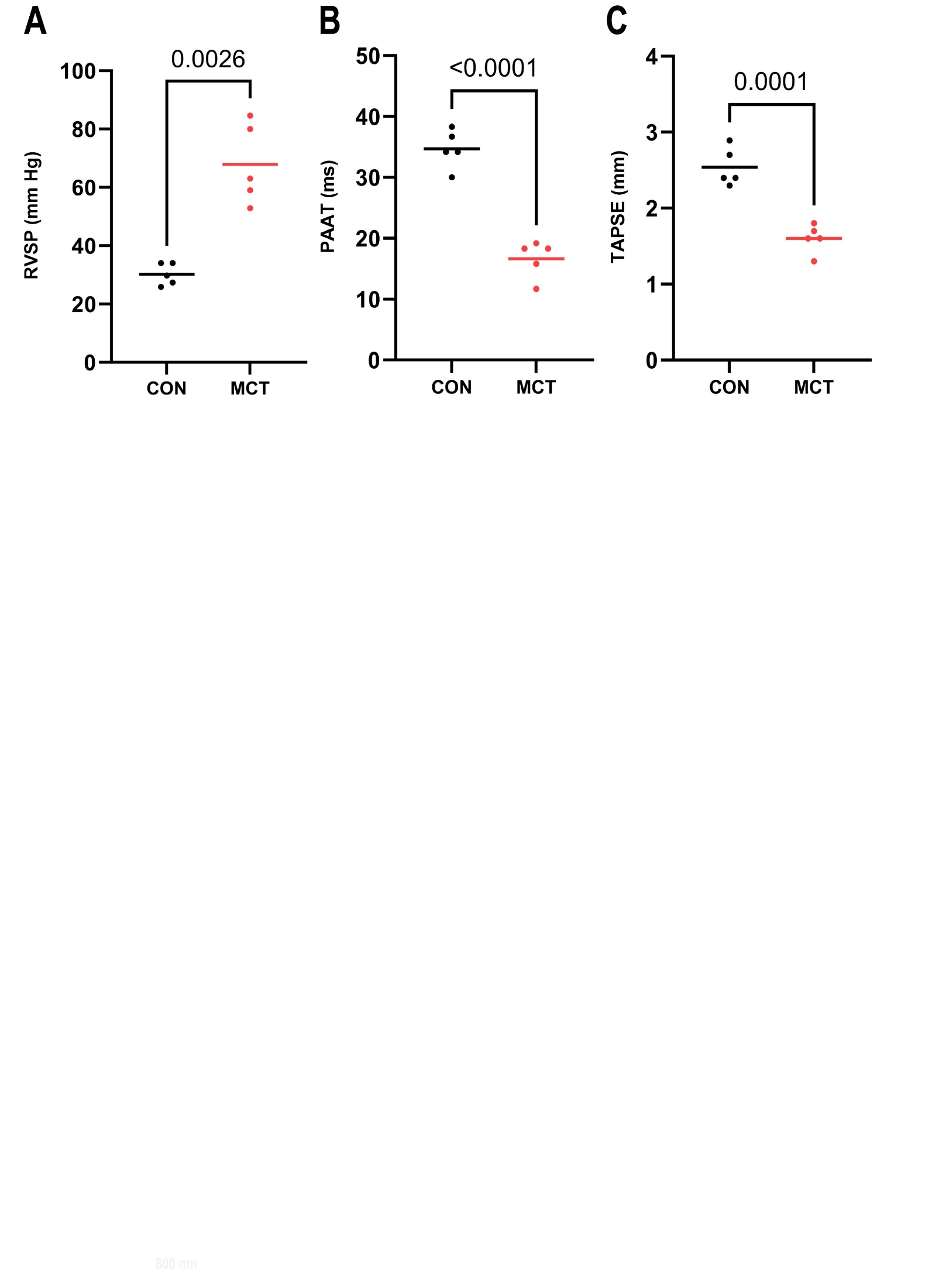
